# Licoisoflavone A improves adipose tissue dysfunction in response to diet induced obesity by promoting METTL3 expression

**DOI:** 10.64898/2025.12.14.694234

**Authors:** Shuangxi Tu, Xiaodan Wang, Haojun Tang, Kai Yin, Xiao Zhu

**Affiliations:** Guangxi Key Laboratory of Diabetic Systems Medicine, Guilin Medical University, No. 1, Zhiyuan Road, Lingui District, Guilin City, Guangxi Zhuang Autonomous Region, China, Postcode 541199; Guangzhou Key Laboratory of Diabetes Metabolic Reprogramming and Precision Prevention and Control, Department of General Practice, The Fifth Affiliated Hospital of Southern Medical University, No. 566, Congcheng Avenue, Conghua District, Guangzhou City, Guangdong Province, China, Postcode 510900

**Keywords:** Obesity, Licoisoflavone A (LIC-A), Methyltransferase 3 (METTL3), adipose tissue(AT), Inflammation

## Abstract

As developing obese adipose tissue, adipocytes suffer pathological changes from inert energy storage to excessive and active endocrine organ in association with disease hazard. METTL3, as an RNA methyltransferase of key importance in adipose tissue development, functional preservation and metabolic homeostasis, is remarkably complex. Licoisoflavone A (LIC-A) is a prenylated flavonoid compound of licorice that has been suggested to display antiinflammatory, -hypertrophic, and -proliferative activities; it remains unknown whether it can directly modulate expression of METTL3 and impact functions of adipose tissue in obesity. Here in our study, 16 candidates in the top ranking molecules were screened out in the process of the molecular docking between our TCM formula Fangjihuangqin Decoction and protein METTL3 (PDB: 5TEY). Among them, LIC-A was considered as a potential upstream regulatory molecule of METTL3 and obviously promote the expression of METTL3. The effects of TNF-α and LPS treatments on the inhibition of adipogenesis were successfully recovered by LIC-A treatment in the adipogenic 3T3-L1 cells, via the regulation of adipogenic cytokines, as well as the expression of inflammatory factors. These protective effects were similarly abolished with METTL3 knockdown, suggesting the role of LIC-A relies on METTL3. Moreover, in vivo data demonstrated that administration of LIC-A could significantly recover the body weight lipid metabolism, insulin resistance and gluconeogenesis in mice with reduced adipose depostion as well as abated systemic inflammation. In conclusion, our study suggested that LIC-A could be the key component in preserving the homeostasis of adipose tissues through regulating METTL3 expression, indicating LIC-A’s promising treatment potential for obesity-related metabolic diseases.

## Introduction

Obesity is a chronic metabolic complex disorder related with a plethora of comorbidities such as metabolic syndrome[1], insulin resistance[2], hypertension, and diabetes[3]. High level of adipose tissue depositing body fat is one of the main features of obesity, accumulated mainly in triglycerides, favoring adipocyte hypertrophy and/or hyperplasia. Dysregulated adipogenic gene expression in hypertrophied adipocytes limits their fat storage capacity and enhanced secretion of inflammatory adipokines. While AT inflammation is usually low grade, continuous stimulation by inflammatory mediators is sufficient to cause major organ dysfunction and leads to onset of metabolic obesity-derived complications[4,5]. Synthetic anti-obesity agents currently at disposal include orlistat, an anti-obesity agent that inhibits gastrointestinal and pancreatic lipase thereby decreasing intestinal fat absorption[6]and liraglutide, an agent that preferentially decreases fat mass but increases lean body mass[7,8]. Pharmacological interventions are an increasingly promising strategy to address obesity. However, concerns on side effects and adverse drug reactions are profound public health issues and a major hurdle in the development of anti-obesity drugs. Consequently, the development of accurate molecular targets for adipose tissue function and targeted treatments is of high relevance.

METTL3 serves as a core catalytic subunit of the m^6^A methyltransferase complex [9–11] and plays a central role in numerous biological processes, including cell proliferation, migration[12,13], and energy metabolism [14–16]. As the most abundant reversible chemical modification in eukaryotic mRNA [17,18], m^6^A methylation is dynamically regulated by “writer” (e.g., METTL3)[19, 20], “eraser” (e.g., FTO)[21], and “reader” proteins (e.g., YTHDF1/2/3)[22–24], which collectively modulate mRNA processing and fate[25,26]. Recent studies indicate that METTL3-mediated m^6^A modification promotes the induction of beige adipocytes within white adipose tissue (WAT), a process associated with improved glucose and lipid metabolism and protection against obesity [27]. Moreover, METTL3 upregulates glycolytic genes via m^6^A modification during cold-induced beige adipocyte activation [28]. These findings highlight the potential of targeting METTL3-mediated m^6^A modification to restore adipose tissue homeostasis.

Obesity drug therapies from natural product offers the additional advantages like fewer side effects of treatment as well as other health benefits like anti-diabetic, anti-hyperlipidemic effects[29, 30] justifying its further exploration. Anti-obesity actions of several traditional herbal formulations was critically evaluated in recent years [31]. There are some reported cases about Fangjihuangqin Decoction, Berberine and fenugreek that indicate their efficacies for modulating obesity and adipose tissue inflammation[32–34] and Fangjihuangqi Tang a formula of Zhang Zhongjing Jin Gui Yao Lue, contains four ingredients Stephaniae tetrandrae Radix (Fangji), Astragali Radix (Huangqi), Atractylodis macrocephalae Rhizoma (Baizhu) and Glycyrrhizae Radix (Gancao). It has been reported that it can enhance the glucose and lipid metabolism, and cause considerable weight loss, becoming the earliest proven traditional anti-obesity formula that had gained the international recognition. However, its active components and the mechanism behind are little known despite its proven efficacy. We therefore speculate that Fangjihuangqi Tang bears bioactive materials capable of regulating METTL3 expression and reversing the disturbance of the adipose tissue. The localization of such compounds may suggest new therapeutic approaches for treating metabolic syndrome associated with obesity. Licoisoflavone A (LIC-A) is one of the principal (natural) isoflavone flavonoids in the root of licorice (Glycyrrhiza uralensis), whose role in obesity has not previously been addressed. Based on molecular docking-based screening of 16 candidate compounds, we selected LIC-A as a candidate, then tested its action on the key genes involved in lipogenesis, lipolysis, and adipose inflammation in TNF-α and LPS stimulated adipogenic 3T3-L1 adipocytes and HFD fed C57BL/6J mice. This makes our data suggesting that LIC-A reduces metabolic dysregulation via boosting METTL3 mediated m^6^A methylation to be promising for its therapeutic relevance for treating obesity.

## Results

### LIC-A monomer in Fangji-Huangqi Decoction promotes METTL3 expression in adipogenic 3T3-L1 cells

In order to screen drug monomers for the enhancing effect in METTL3 expression and adipose tissue homeostasis, we firstly screened Traditional Chinese medicine formula Fangjihuangqin Decoction through the Traditional Chinese Medicine Systems Pharmacology Database and Analysis Platform(TCMSP), and then subjected monomeric compounds to molecular docking simulations with protein METTL3 (PDB: 5TEY) using Discovery Studio. We considered compound–protein interactions and executed molecular force field calculations for evaluating binding affinity and biological activity. We used LibDock to sort docking scores into binding free energy and molecular affinity(Supplementary Table 1) to select 16 drug-like monomers (Supplementary Figure 1A).

We first determined the optimal non-cytotoxic treatment concentration for each monomer in adipogenic 3T3-L1 cells using MTT assays across a range of concentrations (Figure 1A). Cells were then treated with each monomer at the selected concentration for 48 hours. Western blot analysis revealed that, among all candidates, LIC-A significantly upregulated METTL3 protein expression compared with other drug monomers and the DMSO control (Figure 1B-C). Moreover, dot blot and colorimetric assays showed that LIC-A treatment markedly increased global RNA m^6^A modification levels (Figure 1D-E). Together, these results indicate that LIC-A enhances METTL3-m^6^A modification in 3T3-L1 preadipocytes.

**Figure 1.**
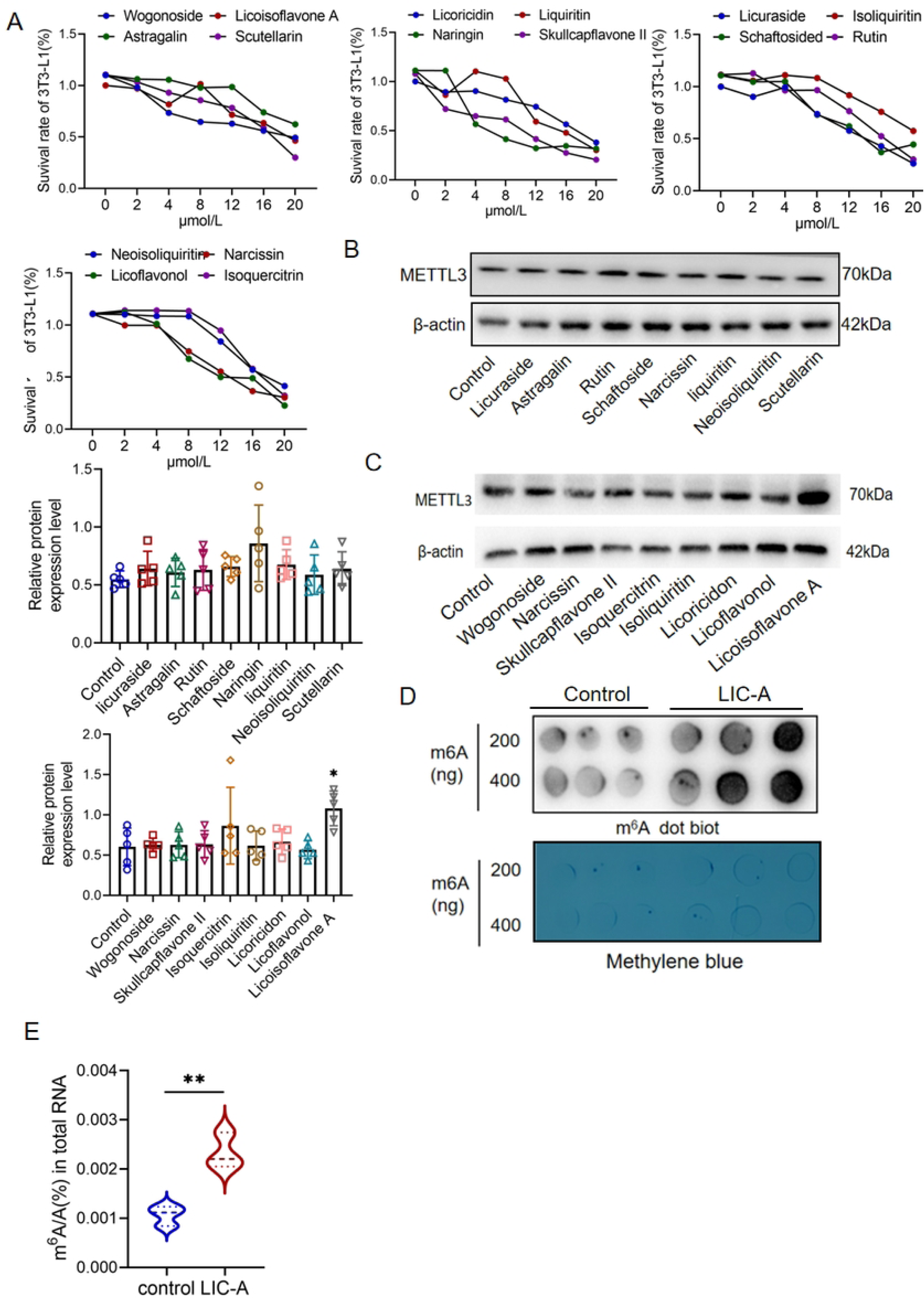
LIC-A monomer in Fangji-Huangqi Decoction promotes METTL3 expression in adipogenic 3T3-L1 cells. **A** Viability of 3T3-L1 cells treated with the indicated drugs at various concentrations, as determined by MTT assay. **B, C** Western blot analysis of METTL3 protein expression in 3T3-L1 cells following treatment with individual drugs. **D** Global m^6^A levels in mRNA from LA-treated 3T3-L1 cells, measured by m^6^A dot blot using an anti-m^6^A antibody. **E** Quantitative analysis of global mRNA m^6^A levels in LA-treated cells using a colorimetric ELISA kit.

### LIC-A ameliorates TNF-α and LPS induced adipokine dysregulation and inflammatory responses in differentiating 3T3-L1 adipocytes

As widely recognized, white adipose tissue acts as an active endocrine and immuneregulating organ that communicates with remote organs such as the brain, liver and muscle through the release of adipokines (e.g., leptin, adiponectin, resistin) and inflammatory factors. To explore the impact of LIC-A on adipocyte functions, we first examined whether it regulates adipokine and inflammatory factor expression in TNF-α or LPS stimulated adipogenic 3T3-L1 adipocytes. We first identified the optimum concentration of LIC-A used under acute TNF-α treatment (Figure 2A). Differentiated adipogenic 3T3-L1 cells were challenged with the obesity-mimicking cocktail and followed by TNF-α treatment. ELISA assay showed that LIC-A treatment could significantly enhance adiponectin secretion and inhibit leukin secretion (leptin and resistin) when compared with group treated by TNF-α alone (Figure 2B-C). LIC-A also inhibited the release of inflammatory cytokines, such as IL-6 and IL-1β, induced by TNF-α (Figure 2D) and elevated the expression of genes of insulin-sensitizing (Figure 2E). At transcription level, LIC-A significantly promoted the expression of adiponectin and leptin mRNA and resistin mRNA reduced compared to the TNF-α group (Figure 2F). In line with, LIC-A decreased both mRNA and protein of TNF-α induced inflammatory factors (Figure 2G-H) significantly.

**Figure 2.**
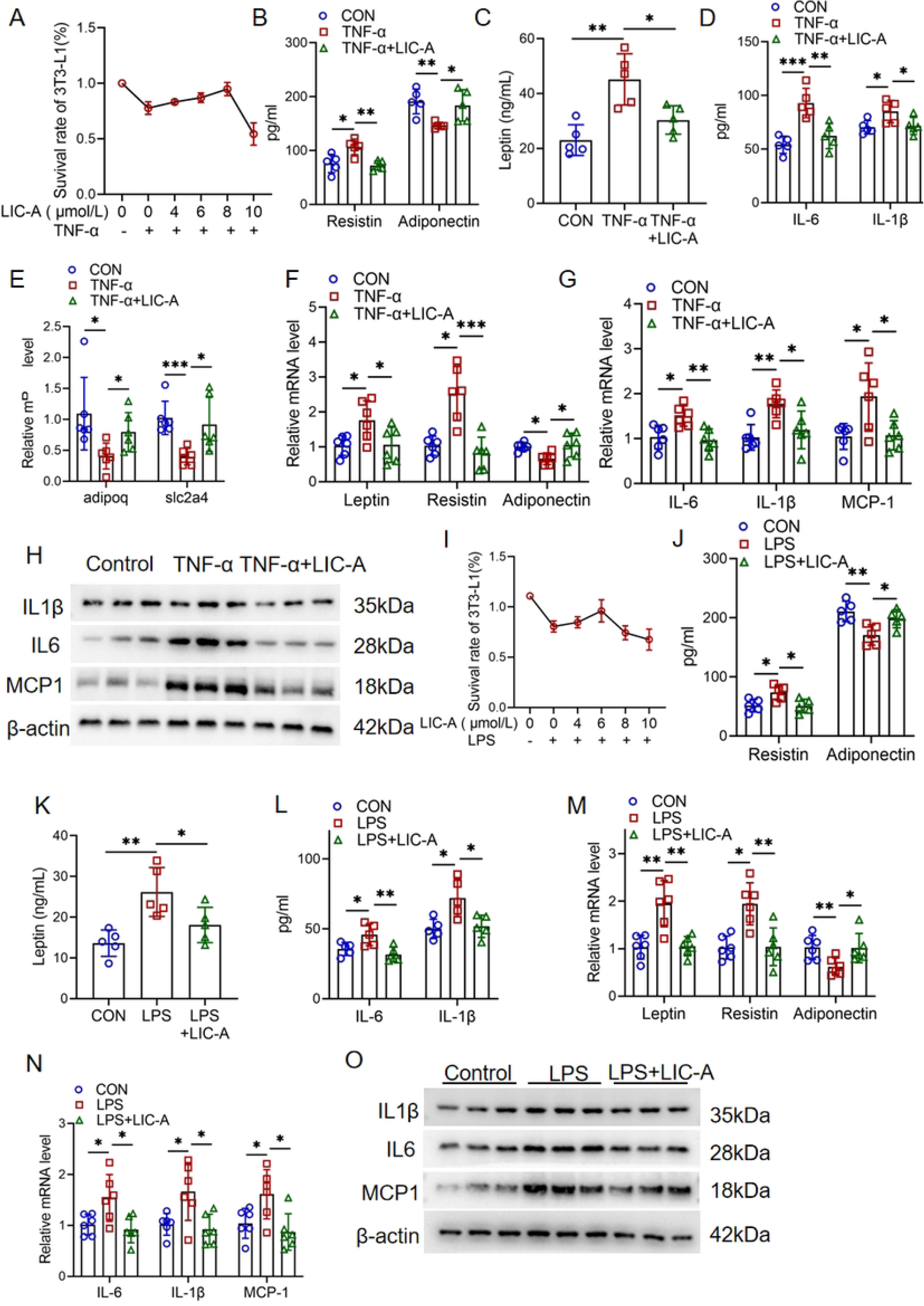
LIC-A ameliorates TNF-α and LPS induced adipokine dysregulation and inflammatory responses in differentiating 3T3-L1 adipocytes. **A** Determining optimal LA concentration under TNF-α stimulation by MTT assay. **B-D** Measuring secreted levels of adipokines and inflammatory cytokines by ELISA. **E** Analyzing mRNA expression of insulin sensitivity genes by qPCR**. F,G** Profiling mRNA expression of adipokines and inflammatory cytokines by qPCR. **H** Detecting protein levels of inflammatory cytokines by Western blot. I Determining optimal LA concentration under LPA stimulation by MTT assay. **J-L** Measuring secreted levels of adipokines and inflammatory cytokines by ELISA. **M,N** Profiling mRNA expression of adipokines and inflammatory cytokines by qPCR. **O** Detecting protein levels of inflammatory cytokines by Western blot.

We further evaluated whether LIC-A exerts similar regulatory effects under acute LPS stimulation. After determining the optimal LIC-A concentration under LPS stimulation (Figure 2I), we observed that LIC-A treatment substantially ameliorated LPS induced alterations in adipokine secretion and inflammatory cytokine production (Figure 2J-L). Moreover, LIC-A regulated mRNA and protein expression of adipokines and inflammatory factors in a manner highly consistent with its effects under TNF-α stimulation (Figure 2M-O). Together, these data indicate that LIC-A alleviates TNF-α and LPS induced dysregulation of adipokine expression and inflammatory responses in adipogenic 3T3-L1 cells.

### METTL3 lacking counteracts LIC-A improved adipokine dysregulation and inflammatory response in TNF-α or LPS stimulated adipogenic 3T3-L1 cells

Since upregulation of METTL3 could be an important element in the improvement of LIC-A by TNF-α and LPS induced adipokine dysregulation and inflammatory responses, we first examine METTL3 protein level in adipogenic 3T3-L1 cells. TNF-α or LPS acute stimulation obviously decreased the METTL3 level, which was again attenuated by LIC-A treatment (Figure 3A). We next transiently transfected adipogenic adipogenic 3T3-L1 cells with a METTL3 knockdown lentivirus, effectively confirming METTL3 knockdown (protein and mRNA) (Figure 3B-C). ELISA revealed that in response to acute TNF-α stimulation METTL3 knockdown abolished the beneficial effect of LIC-A. This was supported by lack of increase in adiponectin secretion, lack of a significant decrease in leptin and resistin expression, and lack of an effect on secretable levels of key proinflammatory cytokines in comparison to the empty vector group (Figure 3D-F). In accord with these secreted changes, METTL3 knockdown blocked the LIC-A upregulation of adiponectin mRNA and the downregulation of leptin, resistin and proinflammatory factor transcripts (Figure 3G-H). We also investigated whether LIC-A could ameliorate secretory dysfunction under LPS challenge in METTL3deficient cells.

**Figure 3.**
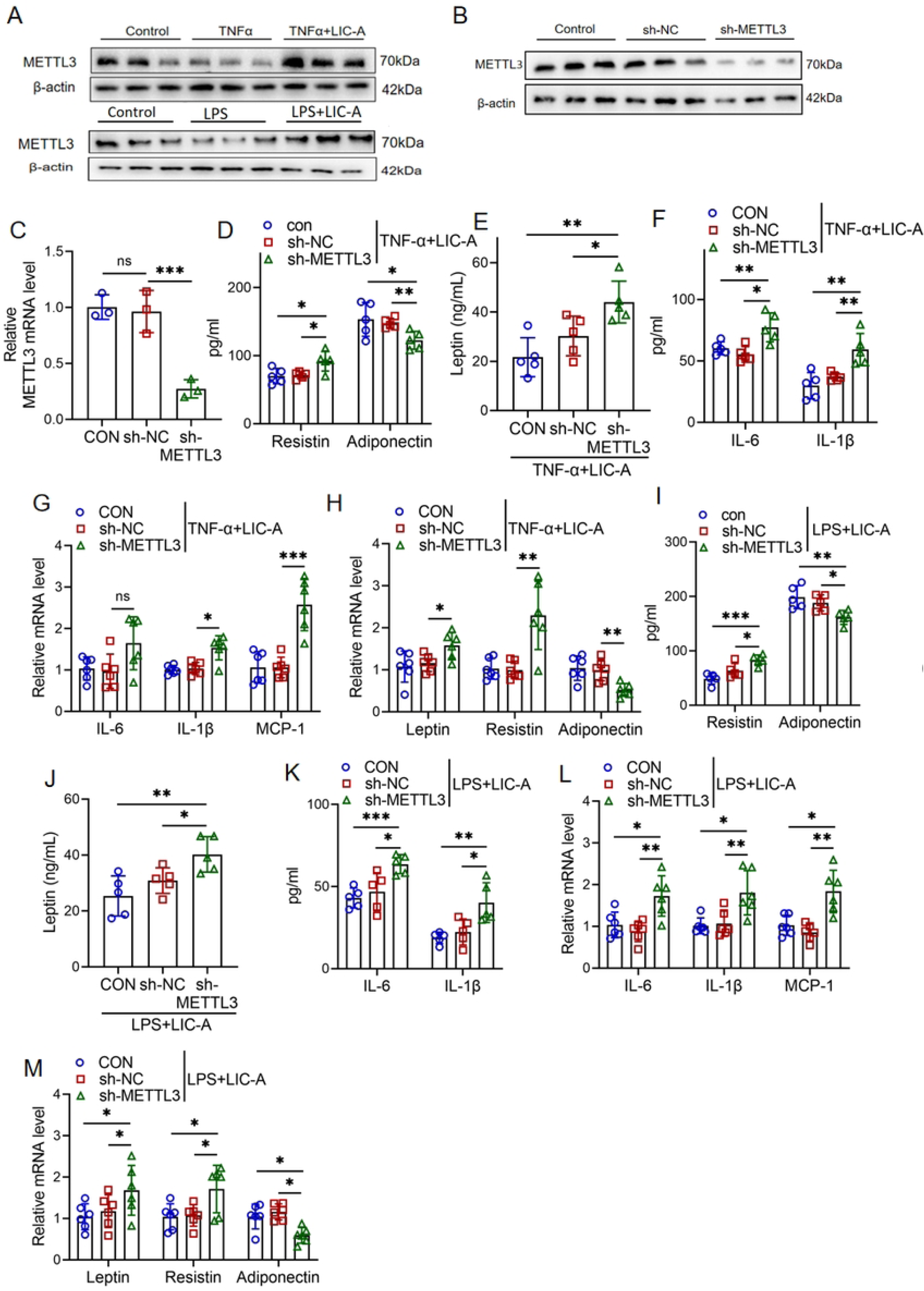
METTL3 lacking counteracts LIC-A improved adipokine dysregulation and inflammatory response in TNF-α or LPS stimulated adipogenic 3T3-L1 cells. **A** Western blot analysis of METTL3 protein expression during adipogenesis in 3T3-L1 cells under acute TNF-α or LPS stimulation. **B** Western blot analysis verifying the knockdown efficiency of METTL3 in adipogenic 3T3-L1 cells. **C** qPCR analysis verifying the knockdown efficiency of METTL3 in adipogenic 3T3-L1 cells. Phenotypic Consequences under TNF-α Stimulation. **D-F** ELISA measuring the secretion levels of resistin,adiponectin, leptin, and inflammatory cytokines. **G, H** qPCR analysis of mRNA levels for resistin adiponectin, leptin, and inflammatory cytokines. Phenotypic Consequences under LPS Stimulation. **I-K** ELISA measuring the secretion levels of resistin,adiponectin, leptin, and inflammatory cytokines. **L,M** qPCR analysis of mRNA levels for resistin adiponectin, leptin, and inflammatory cytokines.

ELISA confirmed that METTL3 knockdown abrogated the protective effects of LIC-A under LPS challenge, as shown by persistently low adiponectin secretion alongside unabated levels of leptin, resistin, and inflammatory mediators. Consistent with this profile, qPCR analysis further revealed a parallel failure in the transcriptional regulation of these key metabolic and inflammatory genes (Figure 3I-M). Collectively, these results indicate that the ability of LIC-A to counteract TNF-α and LPS induced adipokine dysregulation and inflammation depends on its promotion of METTL3 expression.

### LIC-A Alleviates HFD Metabolic Disorder and Adipose Tissue dysfunction in Obese Mice

To assess the in vivo anti-obesity effects of LIC-A, obesity was induced in C57BL/6J mice with high-fat diet (HFD) over 12-week period. The experimental animal (from 4 to 16 weeks) were injected LIC-A (30 mg/kg) by oral gavage (twice a week) (Figure 4A). The body weight of the experimental group treated LIC-A decreased remarkably in comparation with that of HFD control group (Figure 4B-C). This metabolic advantage was associated with improved insulin sensitivity and glucose homeostasis (Supplementary Figure 4A-B) and strikingly with reduced hyperglycemia and dyslipidemia (Supplementary Figure 4C). In addition, LIC-A efficiently attenuated WAT mass enlargement, and impeded pathological re-modeling of this tissue (Figure 4D-E). We also noticed considerable reduction in size of adipocytes in eWAT of LIC-A treated animals (Figure 4F-G and Supplementary Figure 4D) that is reflection of inhibition of hypertrophy of adipocytes which is a hall mark of dys-functional adipose tissue in obesity.

**Figure 4.**
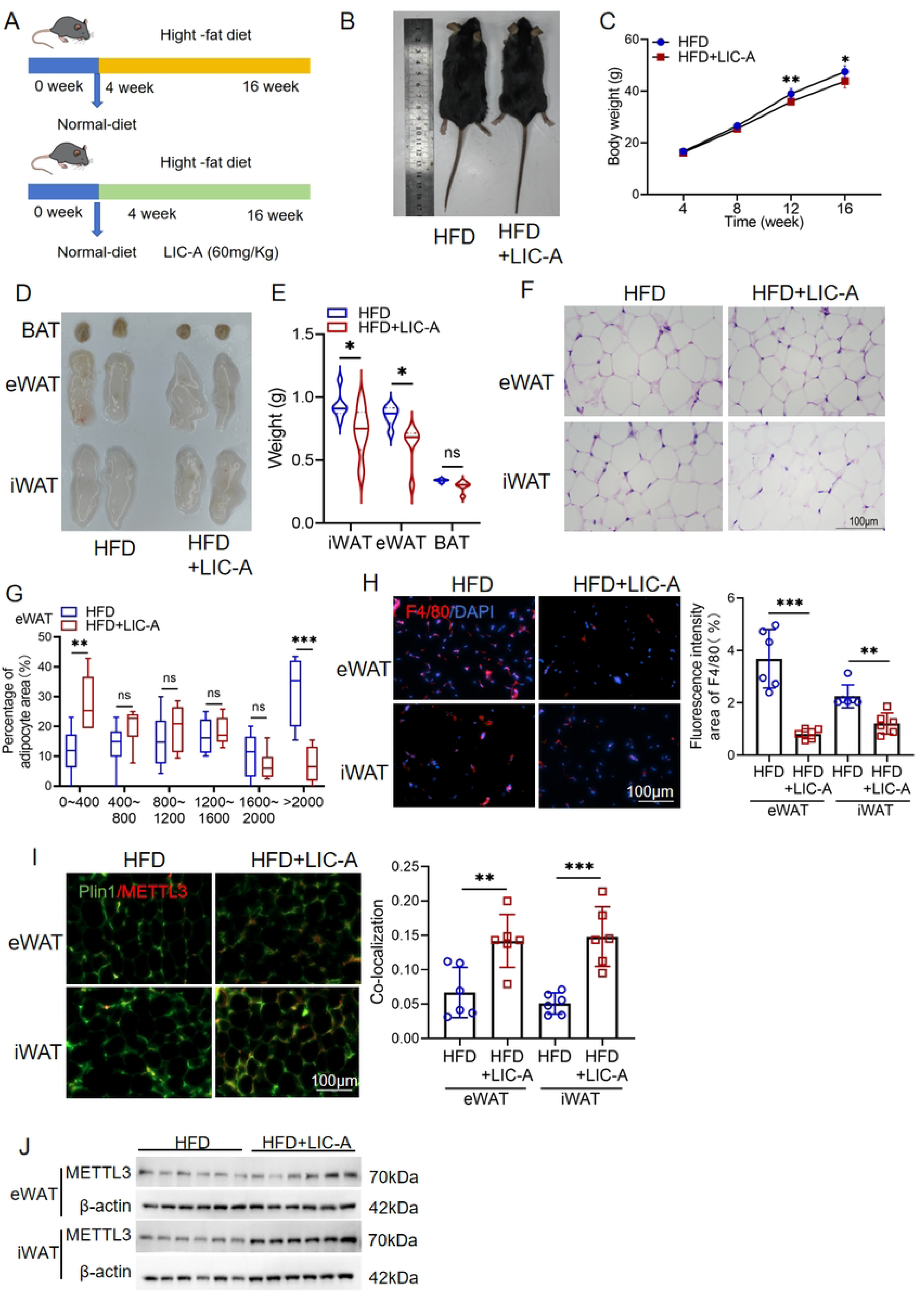
LIC-A Alleviates HFD Metabolic Disorder and Adipose Tissue dysfunction in Obese Mice. **A** Schematic of the HFD feeding experimental strategy. **B** Representative morphology of mice after 16 weeks. **C** Monitoring of body weight changes. **D, E** Examination of adipose tissue morphology and weight. **F** Histological assessment of adipose tissue by H&E staining. **G** Quantification of adipocyte size frequency distribution in eWAT. **H** Assessing macrophage infiltration by F4/80/DAPI immunofluorescence and quantitative analysis of inflammatory areas. **I** Analyzing METTL3 localization by Plin1/METTL3 immunofluorescence and quantitative analysis of METTL3-positive areas. **J** Detecting METTL3 protein levels in WAT by Western blot.

Notably, in the presence of LIC-A adipose tissue inflammation was reduced, since inflammation markers measured as macrophage infiltration was diminished (Figure 4I). Indeed, consistently the whole panel of adipokines evidenced a metabolically favorable shift. In line with this, ELISA using adipose tissue homogenates showed increased adiponectin and decreased levels of leptin, resistin and inflammatory factors (at protein level). Furthermore, qPCR analysis of inflammatory gene expression at the transcript level confirmed the downregulation of proinflammatory genes (Supplementary Figure 4E-H).

Notably, we found that LIC-A treatment also elevated METTL3 expression in the AT of mice, thereby recapitulating our in vitro findings and confirming that LIC-A promotes METTL3 expression in both physiological and experimental settings. (Figures 4I-J). Taken together, these results demonstrate that LIC-A supplementation effectively alleviates metabolic dysregulation and adipose tissue harmful changes in diet-induced obese mice, suggesting that LIC-A’s beneficial effects are mediated, at least in part, through METTL3 upregulation in adipose tissue.

## Discussion

Obesity and the comorbid conditions, known as metabolic diseases, are associated with dysregulated adipose tissue that becomes pathologically remodeled and chronically inflamed and secretes adipokines in an altered fashion[35–37]. Here, we pinpointed the licorice isoflavone A (LIC-A) as a new upstream controller of the RNA methyltransferase METTL3 and showed that LIC-A alleviates the metabolic disturbances of obesity by stimulating the METTL3mediated m^6^A RNA methylation in the adipose tissue.

In summary, our data indicate for the first time that the natural product LIC-A (isolated from licorice, a traditional Chinese medicine), upregulates the expression of the METTL3, in adipogenic 3T3-L1 adipocytes, and is accompanied by a global increase in the amount of modification, m^6^A, suggesting an involvement of LIC-A effects on their metabolism through an epitranscriptomic mechanism. Significantly, LIC-A treatment reversed TNF-α and LPS caused deregulation of adipokines (leptin, adiponectin and resistin) and inhibited production of the pro-inflammatory cytokines. This was entirely lost upon METTL3 knockdown and this establishes an etiological link between LIC-A’s anti inflammatory and metabolic effects and METTL3 expression.

In vivo, LIC-A was effective in strongly improving systemic glucose homeostasis, insulin sensitivity and lipid profiles in HFD-fed C57BL/6J mice, which was associated with decreased adipocyte hypertrophy and macrophage infiltration, and ameliorated inflammation in eWAT. Similar to our cellular data, LIC-A resulted in higher METTL3 protein levels in eWAT in further evidence that METTL3 is an important mediator of the pro-beneficial effect of LIC-A.

Our findings align to the accumulating evidence linking METTL3 to be the key regulator of plasticity of adipose tissue and metabolic health[38]. Previous studies have demonstrated that METTL3 participates in beige adipogenesis and enhances glycolytic gene expression through m6A modification, thereby promoting energy expenditure and glucose tolerance[27,28]. Here we provide the basis for such a therapeutic approach, by finding a natural compound able to potentiate a positive modulation of the METTL3 activity.

Significantly, adipose endocrine dysfunction is another mechanism that possibly account for antiobesity actions of LIC-A. Since restoration of adipokine homeostasis and combating adipose tissue inflammation, may explain, at least partially the maintenance of a state of cellular non-dysfunction that could compensate not only to reduce insulin resistance and dyslipidemia that are the physiological signs of the systemic effects of adipose tissue dysfunction. In this regard the decreased levels of leptin and resistin levels, accompanied by increased adiponectin suggests that the improvements of adipose endocrine dysfunction that the intake of LIC-A caused, are essential to the state of metabolic homeostasis.

The following caveats of our study merit to be mentioned. First, we identified LIC-A to be a METTL3 enhancer by molecular docking and functionally, and the direct binding event of LIC-A with METTL3 has yet to be proven by biophysical means (e.g., surface plasmon resonance, crystal structure). Second, the downstream targets of METTL3 that mediate the LIC-A induced improvement in metabolism needs to be explored further. RNA sequencing and m^6^A-seq could detect the individual transcripts that are whose methylation and expression are regulated by LIC-A. Third, the translational significance of our data should be tested in human adipocytes or clinical context.

Overall, in this work we confirmed LIC-A as a first-in-class, natural agonist of METTL3 which is extremely promising therapeutically for both obesity and metabolic disorder. Reconciling LIC-A with METTL3-catalyzed m^6^A methylation offers a mechanistic basis for how traditional Chinese medicine compounds can influence epigenomic composition and reverse metabolic imbalances. Additional research will include improving LIC-A based treatments and testing the clinical potential of this approach in patient models for disease.

## Competing Interest Statement

The authors declare no competing interests.

## Acknowledgements

This study was supported by the National Natural Sciences Foundation of China (grant numbers 82370463), Natural Science Foundation of the Guangxi Zhuang Autonomous Region (grant numbers 2024GXNSFAA010154), Guangdong Basic and Applied Basic Research Foundation (grant numbers 2025A1515012522), the China Postdoctoral Science Foundation under Grant Number 2024M761313.

## Materials and methods

### Animal Experiments

The animal study was conducted under a protocol approved by the Institutional Animal Care and Use Committee of Guilin Medical University, adhering to the National Institutes of Health guidelines for laboratory animal welfare. Female C57BL/6J mice (Cyagen Biosciences Inc.) were acclimatized under standard SPF conditions with a 12-hour light/dark cycle, temperature maintained at 24 ± 2°C, and humidity at 50 ± 10%. Mice were fed a high-fat diet (HD001, 60% KCal, Biotech-HD Co., Ltd) ad libitum. Following a 1-week acclimatization, mice were randomly assigned using a computer-generated randomization scheme to two groups (n= X per group): the HFD control group (vehicle) and the LIC-A treatment group (30 mg/kg). Treatments were administered by oral gavage twice weekly from week 4 to week 16 post-diet initiation.

### Molecular Docking

We applied molecular docking to screen METTL3 binder candidates in Fangji-Huangqi Decoction. Initially we construct a compound library by mining chemical components in 4 main herbs of Fangji-Huangqi Decoction, i.e., Astragalus membranaceus, Stephania tetrandra, Scutellaria baicalensis, Glycyrrhiza uralensis, using the TCMSP database. We filtered this library by drug-likeness, leaving 528 compounds. After that, the structure of METTL3 (PDB: 5TEY) was prepared in Discovery Studio 2021 by deleting the water, adding the hydrogens and assigning charges. Then, flexible docking was run from the LibDock module with defining the binding pocket by using a 5 Å radius, around the native ligand. Ligand - protein affinities were finally ranked as per the LibDockScore.

### Cell Culture

3T3-L1 cells The cells were maintained in DMEM with 10% heat-inactivated foetal bovine serum (Sangon Biotech Shanghai Co., Ltd.) with 1% penicillin and streptomycin procured from Sigma (St. Louis, MO, USA) in a humidified environment with 5% CO2 at a constant temperature of 37 °C.

### Cocktail Induction Method

To initiate adipogenic differentiation, 3T3-L1 cells on T25 flasks or 6-well plates were allowed to attain confluency and fresh medium (standard) was replaced with adipogenic differentiation medium including 0.5 mM IBMX, 1 uM dexamethasone and 1 ug/mL insulin. Three days following induction, the differentiation medium was replaced with DMEM (containing 10% FBS and 1 u g/mL insulin) as the maintenance medium. Every two days, a fresh volume of maintenance medium was renewed to ensure differentiation until day 8.

### MTT detection of cell viability

3T3-L1 cell were incubated with 5 mg/mL MTT solution (MedchemExpress, HY-15924) for 4 h. Dimethylsulfoxide (DMSO) was added to each well, and the plate was shaken for 10 min to completely dissolve the crystals. The absorbance of each well was measured using a microplate reader (optical density measured at 490 nm). Finally, cell viability was calculated.

### RNA m^6^A quantitative assay

Relative m^6^A levels among total RNAs were evaluated using a m^6^A RNA Methylation Quantification Kit (Colorimetric). Total RNA was extracted using TRIzol extraction reagent, strips were treated with 200 µg/μl total RNA. According to the instructions, RNA was allowed to bind to the strip upon incubation in high binding solution at 37 °C for 90 min. Next, capture and detection antibodies were added to detect the m^6^A colorimetric levels by reading the absorbance at a wavelength of 450 nm. The relative absorbance was quantified for three experimental replicates for each reaction.

### m^6^A dot blot assays

Total RNA was harvested as previously described with RNA quantity monitoring. RNA was denatured by heating at 95 °C for 10 min and then preserved on ice. Then, mRNAs were blotted onto an N+ nylon membrane. Afterwards, the membrane was treated with ultraviolet cross-linking and blocked with 5% nonfat milk in PBST for one hour at room temperature. The membranes were incubated with a m^6^A-specific antibody (1:2000, Proteintech) for reactivation at 4 °C overnight. After washing, the membranes were incubated with a goat anti-rabbit IgG HRP antibody (1:5000) with gentle shaking for 1 h at room temperature. After washing, the membranes were incubated with an ECL detection reagent and visualized using the detection system.

### ELISA for IL-6, IL-1β、adiponectin, resistin and leptin

The concentrations of IL-6, IL-1β, adiponectin, resistin and leptin in the cell culture supernatants of 3T3L-1 cell supplemented with LIC-A were measured using IL-6, IL-1β, adiponectin, resistin and leptin ELISA kits (UpingBio Technology Co., Ltd, cn). The ELISA was performed according to the manufacturer’s instructions, and the sample readings were obtained at 450 nm using a microplate reader. All samples were assayed in triplicate.

### Quantitative real-time polymerase chain reaction (qRT-PCR)

Total RNA was extracted from HUVECs using TRI reagent (Invitrogen Corporation) according to the instructions of the manufacturer. cDNA was synthesised via reverse transcription, and mRNA levels were examined using 2xTaq PCR Master Mix (Wuhan Huaruikang Biologic Technology Co., Ltd.). The results were normalised with β-actin, and qRT-PCR was performed using real-time fluorescent quantitative PCR (Thermo Fisher Scientific, USA). The primer sequences are listed in SupplementaryTable 2.

### Western blotting

Total protein was extracted from the 3T3L-1 cells or adipose tissue using precooled RIPA buffer (Beijing Solarbio Science & Technology Co., Ltd.). After protein quantification, an equivalentprotein sample was added, separated using 10% sodium dodecyl sulfate-polyacrylamide gel electrophoresis (SDSPAGE), and transferred to 0.45 μL polyvinylidene fluoride (Merck Millipore, Germany). After sealing with 5% skim milk from TBST for 1.5 h, the membrane was incubated at 4 °C 10–12 h with the corresponding primary antibody METTL3 (1:10000, Proteintech), MCP-1 (1;2000, Proteintech, IL6(1:2000, Proteintech),, and β-actin (1:5000, Proteintech). After 12 h, the polyvinylidene difluoride (PVDF) membranes were washed thrice with TBST and incubated for 2 h at 4 °C with their respective secondary antibodies: horseradish peroxidase-labelled goat anti-mouse and goat anti-rabbit IgG (1:4000, Sam Golden Bridge Biotechnology Co.,Ltd, CHN). Finally, the PVDF membranes were imaged using a Molecular Imager ChemiDoc XRS+system (BioRAD, USA). The protein expressions were quantified using ImageJ software and normalised using β-actin as an internal reference.

### Immunofluorescence

Mouse ATs were fixed with 4% paraformaldehyde and permeabilised with 0.5% Triton X-100 in PBS, followed by overnight blocking with 2% PBA (PBS containing 2% bovine serum albumin) and incubated overnight with the appropriate primary antibody METTL3 (1:200, Proteintech), F4/80 (1:200, Proteintech) and Plin1(1:300, Proteintech). A fluorescent-labelled secondary antibody (1:4000, Proteintech) was then added and incubated at room temperature for 1h. After washing with PBS, the nuclei were stained with an antifade mounting medium containing DAPI (Beijing Solarbio Science & Technology Co., Ltd, Beijing, China). Finally, immunofluorescence was examined using confocal microscopy. DAPI was used to counter-stain the nuclei and images were collected by fluorescence microscope.

### Lentiviral Transduction

To establish METTL3-knockdown 3T3-L1 cell models, lentiviral transduction was performed. Briefly, when cells reached approximately 60% confluence in culture plates, they were transduced with METTL3-knockdown lentivirus (titer: 3.5×10^8^ TU/mL) or a corresponding negative control virus. The volume of viral supernatant used was calculated based on the predetermined optimal multiplicity of infection (MOI), the viral titer, and the number of cells. Following a 16-hour incubation period, the virus-containing medium was replaced with fresh complete medium. To select for stably transduced cells, culture was continued in medium supplemented with 5 μg/mL puromycin for 2 days. Subsequently, the resistant cell populations were used for follow-up experiments.

### ITT and GTT

At the 16-week time point, metabolic phenotyping of the mice was performed via glucose and insulin tolerance tests. For the glucose tolerance test (GTT), mice were subjected to a 12-hour fast, followed by an intraperitoneal injection of D-glucose (2 g/kg body weight) prepared as a 400 mg/mL solution. Blood glucose levels were measured from tail vein blood using a glucose meter (Kefu, Qingdao, China) at 0, 15, 30, 60, 90, and 120 minutes post-injection.

The insulin tolerance test (ITT) was conducted after a 3-day recovery period following the GTT. Mice were fasted for 4 hours and then administered an intraperitoneal injection of insulin at a dose of 0.75 U/kg body weight. Blood glucose concentrations were similarly monitored at 0, 15, 30, 60, 90, and 120 minutes after insulin administration. All glucose measurements were recorded and subjected to subsequent statistical analysis.

### Statistical analyses

All experimental data in this experiment are represented as the mean±standard deviation (SD). Comparisons between multiple groups were performed using one-way analysis of variance (ANOVA). Data analysis was performed using GraphPad software, and p<0.05 was considered as a significant difference.

---

**Supplementary Figure1.**
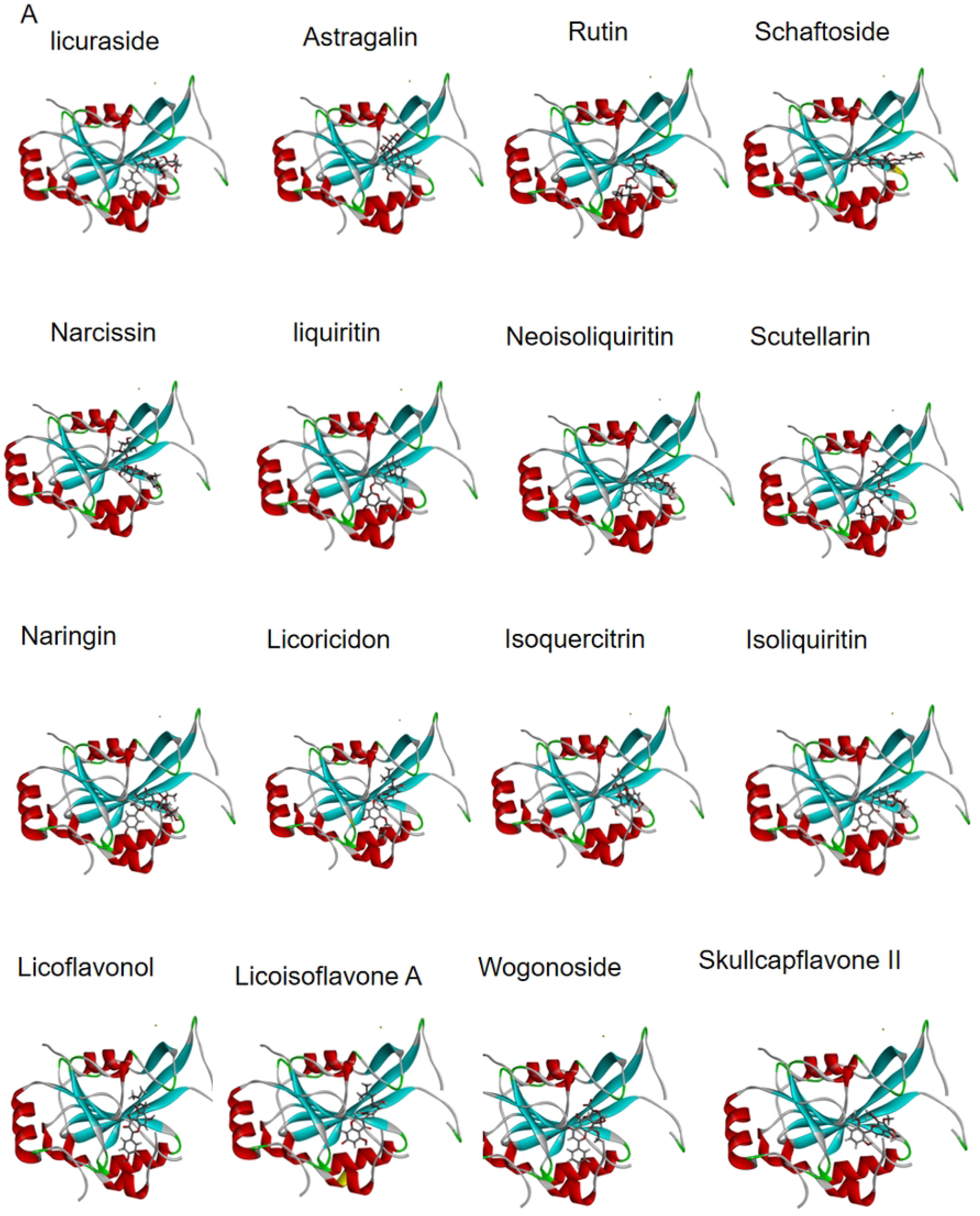
**A** Schematic representation of the molecular docking between the METTL3 protein and drug monomers, performed using Discovery Studio software.

**Supplementary Figure 4.**
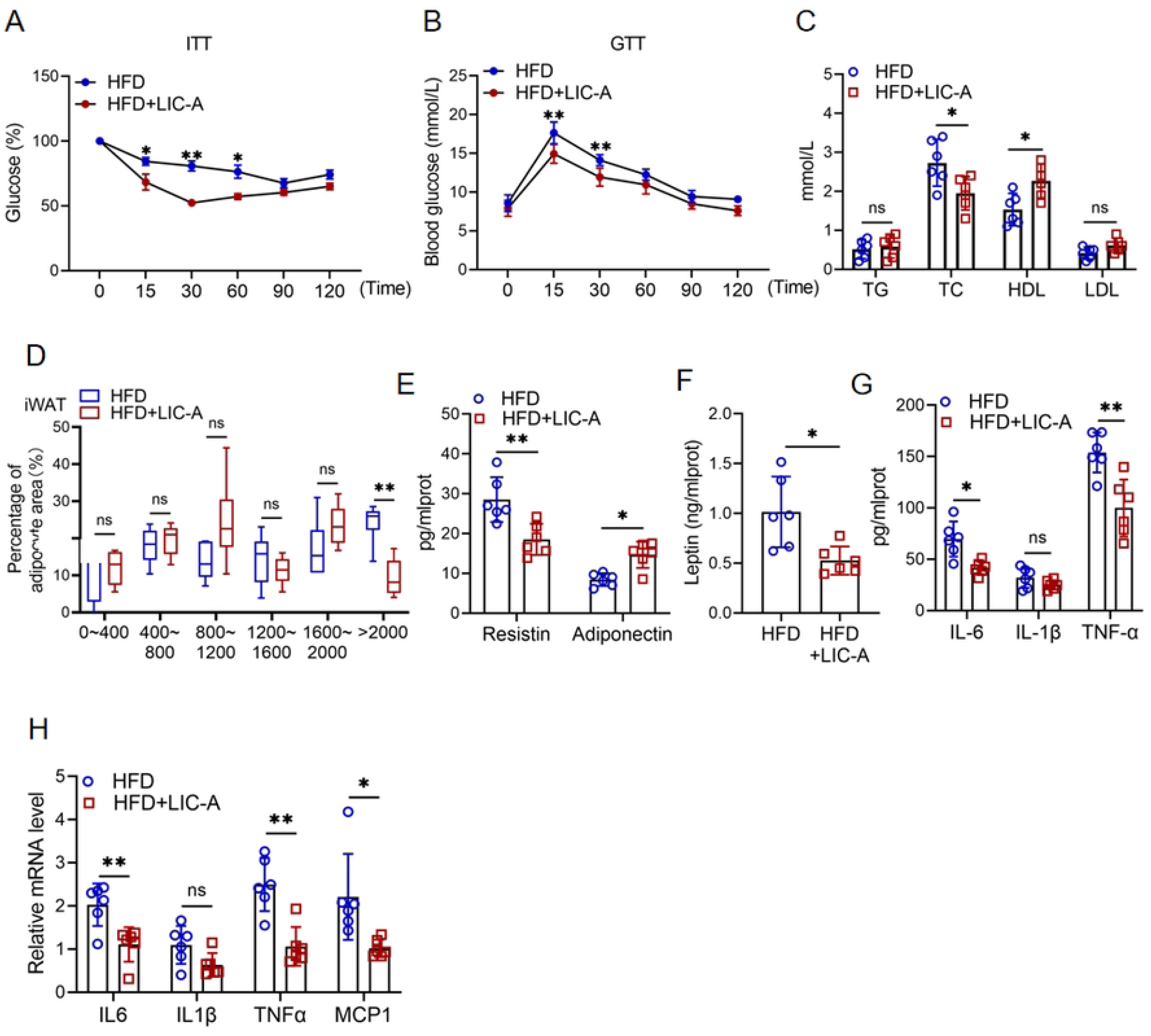
LIC-A Alleviates HFD Metabolic Disorder and Adipose Tissue dysfunction in Obese Mice. **A, B** Valuating insulin and glucose tolerance by ITT and GTT. **C** Measuring blood lipid levels. **D** Quantifying adipocyte size frequency distribution in iWAT. **E-G** Determining serum levels of adipokines and inflammatory cytokines by ELISA**. H** Profiling mRNA levels of inflammatory cytokines by qPCR.

## Notes

### Competing Interest Statement

Declaration of competing interests All authors have reviewed the final version of the manuscript and given approval for its publication. The authors have no conflicts of interest to declare.

